# Soil microbe-induced plant resistance alters aphid inter-genotypic competition leading to rapid evolution with consequences for plant growth and aphid abundance

**DOI:** 10.1101/2022.05.05.490657

**Authors:** Xinqiang Xi, Sharon E. Zytynska

## Abstract

1. Plants and insect herbivores are two of the most diverse multicellular groups in the world, and both are strongly influenced by interactions with the belowground soil microbiome. Effects of reciprocal rapid evolution on ecological interactions between herbivores and plants have been repeatedly demonstrated, but it is unknown if (and how) the soil microbiome could mediate these eco-evolutionary processes.
2. We tested the role of a plant-beneficial soil bacterium (*Acidovorax radicis*) in altering eco-evolutionary interactions between sap-feeding aphid herbivores (*Sitobion avenae*) feeding on barley (*Hordeum vulgare*). We reared two aphid genotypes separately or together on three barley varieties that were inoculated with or without *A. radicis* bacteria. In the first experiment we counted the aphid number and plant biomass after 7, 14 and 21 days of aphid growth, while in a second experiment we counted and removed offspring every 1-2 days to assess aphid longevity and fecundity.
3. Results showed that *A. radicis* increased plant growth and suppressed aphids of both genotypes. The strength of effect was dependent on aphid genotype and barley variety, while the direction of effect was altered by aphid population mixture. Fescue aphids experienced increased growth when they were sharing the plant with Sickte aphids on inoculated plants; this increase was not seen in the control plants without *A. radicis* and was only apparent after 14 days of aphid population growth.
4. Plant inoculation with *A. radicis* reduced aphid survival (reduced number of reproductive days) and fecundity (reduced daily reproductive output for surviving aphids). In the second experiment, when density was controlled, Fescue aphids did not experience increased reproduction in mixed populations, suggesting this is a density-dependent effect. Using Lotka-Volterra modelling, we demonstrated that *A. radicis* inoculation decreased aphid population stability as it increased inter-genotype competition but decreased the intra-genotype competition (likely through reduced population density).
5. Our work demonstrates the important role that plant-associated microbiomes can have in mediating eco-evolutionary interactions between herbivores and host plants.

## Introduction

Defence and counter-defence between plants and herbivores are considered to be one of the main forces to produce and maintain their diversity (Ehrlich & Raven 1964). Rapid evolution of herbivorous insects occurs when genotype frequencies of the same species changes in response to altered biotic and abiotic environmental factors (Hairston *et al*. 2005; Lankau 2011). As different herbivore genotypes always differ in intrinsic growth rate and other ecological characters, rapid evolution may, to some extent, change the species population growth rates (Turcotte, Reznick & Hare 2011). Rapid evolution may also alter predator-prey interactions in a relatively short time period (Yoshida *et al*. 2003). For example, increase in the proportion of higher resistance genotypes in drosophila populations can result in lower parasitism rate and the decreased parasitoid abundance (Zareh, Westoby & Pimentel 1980). In addition, rapid evolution induced population growth (faster or slower) in herbivorous insects may further feedback to alter plant growth (Agrawal *et al*. 2012). Ecologists tend to think evolution occurs much slower compared to ecological processes, thus the occurrence and consequences of rapid evolution has received less attention. It has now been well-recognized that ecological and evolutionary processes can occur at the same timescale (Thompson 1998; Hairston *et al*. 2005; Schoener 2011; Rudman *et al*. 2022). However, mechanisms and consequences of herbivore rapid evolution are still largely unknown (Des Roches *et al*. 2018).

Variation in plant susceptibility to herbivores is a main factor that alters herbivore genotype frequency and the rapid evolution in both direct and indirect ways (Futuyma & Agrawal 2009). Firstly, different herbivore genotypes may suffer in varied extents to induced plant defence and some genotypes may gain higher fitness advantage than others if they employ effective ways to counter the evolved plant defences (War *et al*. 2018). In addition, changes in plant nutrient status along with the varied plant defence may increase or decrease the competition strength and then shift the competition outcome between interacting herbivore genotypes (Turcotte, Reznick & Hare 2013). Secondly, varied plant defence may shift the outcome of plant-mediated inter-genotype interactions, and consequently the genotype frequency. Abundant evidence has clearly demonstrated that changed plant susceptibility can trigger plant-herbivore coevolution, ever since Ehrlich & Raven raised the concept of coevolution (Ehrlich & Raven 1964; Janz 2011). However, if and how altered plant susceptibility due to their interactions with other living beings could induce rapid evolution in herbivore insects are still largely unknown (Howard, Kalske & Kessler 2018; Petipas, Geber & Lau 2021).

Plants naturally recruit and establish associations with a variety of soil microbes that aid the plants in rapidly changing abiotic or biotic conditions (reviewed in Pozo *et al*. 2021; Zytynska 2021). Changes in a host’s microbiome has the potential to shift a host’s phenotype and change the variance in host phenotype in the population, which could shape a host’s response to selection (Henry *et al*. 2021). Inoculating plants with specific bacteria can induce plant defences that reduce the growth rate of feeding herbivorous insects. For example, inoculating plant roots with *Bacillus* spp. can reduce whitefly on cauliflower (Al Arabiat *et al*. 2018), fall armyworm on bermudagrass (Murphey Coy, Held & Kloepper 2017) and aphids on cabbage (Gadhave *et al*. 2016), wheat (Veselova *et al*. 2019) and *Arabidopsis* (Harun-Or-Rashid *et al*. 2017). Termed microbe-induced resistance (MiR; Zehnder *et al*. (1999)), this research is increasingly recognised for its strong potential for sustainable agriculture (Trivedi *et al*. 2021). While the aim of such applied research would be to identify strains of bacteria with general effects across multiple plant and insect species and genotypes, this is rarely the case. Recent work aiming to understand the mechanisms of effect on insects has observed large variation in effect sizes across plant genotypes (Zytynska *et al*. 2020; Wehner *et al*. 2021) and insect phenotypes (Serteyn *et al*. 2020). Such variation has the potential to alter insect population dynamics with evolutionary consequences if changes persist over time.

Aphids are sap-sucking insects that exhibit substantial variation in acceptance and subsequent growth rates across different plant genotypes (Ninkovic & Åhman 2009). As aphids feed they induce plant defences, which also vary dependent on the plant (Delp *et al*. 2009) and aphid genotype (Zytynska *et al*. 2016). In response, aphids can deliver effector proteins to the plant via their saliva that suppress plant defences, enabling higher reproductive output for the feeding aphids particularly in the local area (Escudero-Martinez *et al*. 2020). In a mixed aphid genotype population, variation in any of these traits among aphid genotypes can lead to an eco-evolutionary feedback loop and rapid evolution in the aphid population (Turcotte, Reznick & Hare 2013). The addition of plant defence-inducing microbes to the system has the potential to alter these traits and interactions, especially if the effect of MiR is stronger for one aphid genotype than the other, and further if these interactions are also dependent on the plant genotype. An extreme outcome would be for microbial inoculation to evolve an aphid population with reduced general response to MiR across multiple plant hosts. However, recent work suggests density-dependent effects where aphid suppression was strongest when plants were under high stress, including biotic (higher aphid densities) or abiotic stress (elevated ozone) (Zytynska *et al*. 2020).

We conducted a set of experiments using an aphid-suppressing bacteria (*Acidovorax radices N35e;* Zytynska *et al*. (2020)) on barley to determine if variation in effects of inoculation across plant and aphid genotypes can affect aphid population dynamics, and if any asynchronous response could lead to rapid evolution in the mixed aphid population. We ask whether (1) *A. radicis* bacteria affects the growth rate of *Sitobion avenae* aphid genotypes over time when reared alone or in mixed populations? (2) if single or mixed aphid populations alter the plant growth promotion effect of *A. radicis*? And, (3) if changes in genotype frequency (rapid evolution) can be used to determine longer-term population dynamics of these aphid populations?

## Materials and Methods

### Study system

Our study species included: (1) three barley (*Hordeum vulgare*) plant varieties: Barbarella (Elsoms Seeds), Chevallier (New Heritage Barley Ltd), Irina (KWS); (2) two genotypes of the English grain aphid *S. avenae* (L.) that have been maintained as low density stock populations on barley variety ‘Chanson’, including a dark brown genotype ‘Fescue’ and a pale pinkish coloured genotype ‘Sickte’; and (3) the rhizobacteria *A. radicis* N35. Bacteria were grown on NB agar plates at 30°C for 3 days and then harvested and suspended in 10mM MgCl_2_ at OD_600_=2.0 (10mM MgCl_2_ was also used for the control treatment with no bacteria).

### Effects of microbe-induced plant resistance and aphid coexistence on plant and aphid growth over time

To explore the effects of soil bacteria on plant growth and defence, we grew three barley varieties (Chevallier, Barbarella and Irina) in soil with and without *A. radicis* inoculation. We then reared two genotypes (Fescue, or Sickte) of aphid (*S. avenae*) alone or together on each barley variety; these aphids have different colour morphs allowing ease of assigning genotype in mixed populations (Fescue are dark brown and Sickte is pink-green). Control plants without aphids were also used to test the effects of soil bacteria on plant growth and damage by aphids. There were 8 replicates for each of the 24 treatments, for a total of 192 pots. We set up the experiment in trays (blocked by bacterial treatment to avoid contamination) and replicates/trays were randomised across the experimental space. The experiment was run in a glasshouse (average temperature of 20°C, with a minimum of 18°C and occasional peaks of higher temperatures depending on the outside weather). At these temperatures’ aphids reproduce asexually and are not impacted from the high temperatures experienced. Therefore, the genotype frequency differences can only be explained by soil bacteria mediated decrease or increased fitness and/or the inter-genotype competition.

We first submerged surface-sterilised seeds assigned for “with Acidovorax” treatment in the bacteria solution for 2 hours or in 10mM MgCl_2_ for “without Acidovorax” treatment. Seeds were germinated in potting soil (Levington’s Advance F1 Compost, low nutrient) and plants grown individually in pots (10cm diameter) and covered with air-permeable cellophane bags to prevent the escape of aphids. Using a fine paint brush, we put two third-instar larvae of each aphid genotype on plants assigned for mixed-genotype treatments, and 4 third-instar aphid larvae on the plants assigned for single aphid genotype treatment. We checked settlement of aphids in the first two days after the beginning of the experiment and replaced aphids that failed to settle on plant leaves or died in the first two days.

We counted the number of each aphid genotype in each pot on day 7, day 14 and day 21 and measured the plant leaf length on day 7, day 14 and day 21. We harvested the plants on day 21, measured the fresh shoot and root biomass after counting the final number of aphids. Dry shoot and root mass were also measured after drying at 65°C for 48 hours.

### Lifetime fecundity and longevity experiment

To explore the effects of *A. radicis* inoculation on aphid life history traits (longevity, fecundity), we conducted an experiment to observe aphid development on each of the barley varieties inoculated with and without *A. radicis*. We grew barley seedlings and introduced third-instar Fescue and Sickte aphids alone or together by the same methods mentioned above.

On the day 5 after beginning of the experiment, we removed all of the nymphs in the morning leaving only 4 newly produced first-instar nymphs on each plant at the afternoon. We counted the number of adult aphids in each pot, and then counted and removed the nymphs they produced every 1-2 days until all adults had died to estimate the aphid lifetime fecundity and longevity (survival).

### Data analysis

All the data analysis was conducted using R v3.6.1 in R Studio v1.2.1335. We used linear models to analyse the effect of bacterial inoculation, barley variety, and aphid treatments on plant shoot and root dry biomass. The first model tested the effect of aphid presence and aphid genotype within presence on plant biomass, while the next model subset the data to include only those plants infested with aphids to test effects of aphid population mixture and aphid genotype. A final model on a further subset of the data that included only single aphid genotype populations was used to test effects of aphid genotype on plant biomass.

We used linear models to test effects of bacterial inoculation, barley variety, and aphid treatments (aphid genotype and population mixture) on aphid numbers at days 7, 14 and 21 (all datasets satisfied model assumptions of normal distribution of residuals). We analysed these effects on total aphid number, and also separately for each aphid genotype. An additional mixed effects model was used to analyse these effects over the whole experimental period on aphid number by genotype with pot as a random effect. Finally, genotype frequency was estimated as the number of Fescue aphids relative to the total number of aphids in each pot, and this was again analysed over the experimental period using a mixed effects model with pot as a random effect; genotype frequency was transformed using the arcsine method for proportional data to satisfy model assumptions.

For the lifetime fecundity and longevity experiment, we used linear mixed effects models to test effects of bacterial inoculation, barley variety, and aphid treatments (aphid genotype and population mixture) on total lifetime fecundity, number of reproductive days and daily reproductive output (per surviving adult), with pot as the random factor; normal distribution of residuals ensured no assumptions were violated by using normal error distribution. Aphid survival was analysed using the Cox proportional hazards regression model, using the R library (survival), as a response to bacterial inoculation, barley variety and aphid treatment (aphid genotype and population mixture).

## Results

### Microbe-induced plant growth promotion effects are altered by aphid population mixture

Inoculation of plants with *A. radicis* (hereinafter *Acidovorax*) increased plant shoot and root biomass (shoot: F_1,179_=36.75, P<0.001; root: F_1,179_=24.48, P<0.001; Fig. 1). Plant biomass was reduced by aphids (shoot: F_1,179_=60.52, P<0.001; root: F_1,179_=10.52, P<0.001), and varied across varieties (shoot: F_1,179_=8.72, P<0.001; root: F_1,179_=10.10, P<0.001) (Fig. 1). For the plants hosting aphids, we found that the strength of response of the plants to bacterial inoculation varied depending on the aphid population (single or mixed genotype), also dependent on the barley variety, only for shoot biomass (*Acidovorax* x barley x aphid population interaction: F_2,119_=3.79; P=0.025, Fig. 1), with no effect on root biomass (F_2,119_=2.22; P=0.113). The effect on shoot biomass was not fully explained by variation in aphid number across these treatments since when we controlled for aphid number, we still detected a significant *Acidovorax* x aphid population interaction on shoot biomass (F_1,131_=3.97; P=0.048). Specifically, the microbe-induced increase in shoot biomass for Barbarella plants was stronger when both aphid genotypes were feeding. Yet, the opposite occurred for Irina plants, where double aphid genotype infestation minimised any microbe-induced plant growth promotion effects. In the subset of single aphid genotype plants, aphid genotype itself did not alter the plant’s response to *Acidovorax* inoculation (*Acidovorax* x aphid genotype interaction: F_1,82_=0.50, P=0.482) showing this is a result specific to the single versus mixed aphid genotype population comparisons. The data suggests a reduced growth promotion effect for Barbarella and Chevallier plants when infested by Sickte, but increased for Irina (Fig. 1), but this interaction was not significant (Barley x *Acidovorax* x aphid genotype: F_2,82_=2.01, P=0.141). Exploring this within the subset of the mixed aphid population data to determine any specific effects, we found that neither the absolute number of aphids at the end of the experiment, day 21 (F_1,17_=0.16, P=0.693) nor the ratio of Fescue and Sickte aphids (F_1,17_=0.10, P=0.752) affected plant biomass. However, there is some evidence that an interaction between the total number of aphids at day 14 and *Acidovorax* treatment, dependent on barley variety affected final plant biomass (F_2,35_=3.88, P=0.030), with no similar interaction involving the ratio of Fescue to Sickte aphids (F_2,35_=0.06, P=0.947). In general, we found low association between aphid numbers and shoot biomass, yet the significant interaction was driven by a change in the direction of this response for *Acidovorax* inoculated Irina and Barbarella plants. Briefly, for Barbarella (with increased *Acidovorax* effect in mixed populations), there was a positive relationship between aphid number and plant biomass (larger plants with more aphids) while no such effect for control plants. While for Irina (with a loss of plant growth promotion) we observed a negative relationship (i.e. larger plants with fewer aphids), with no similar effect observed for control plants.

**Figure 1.**
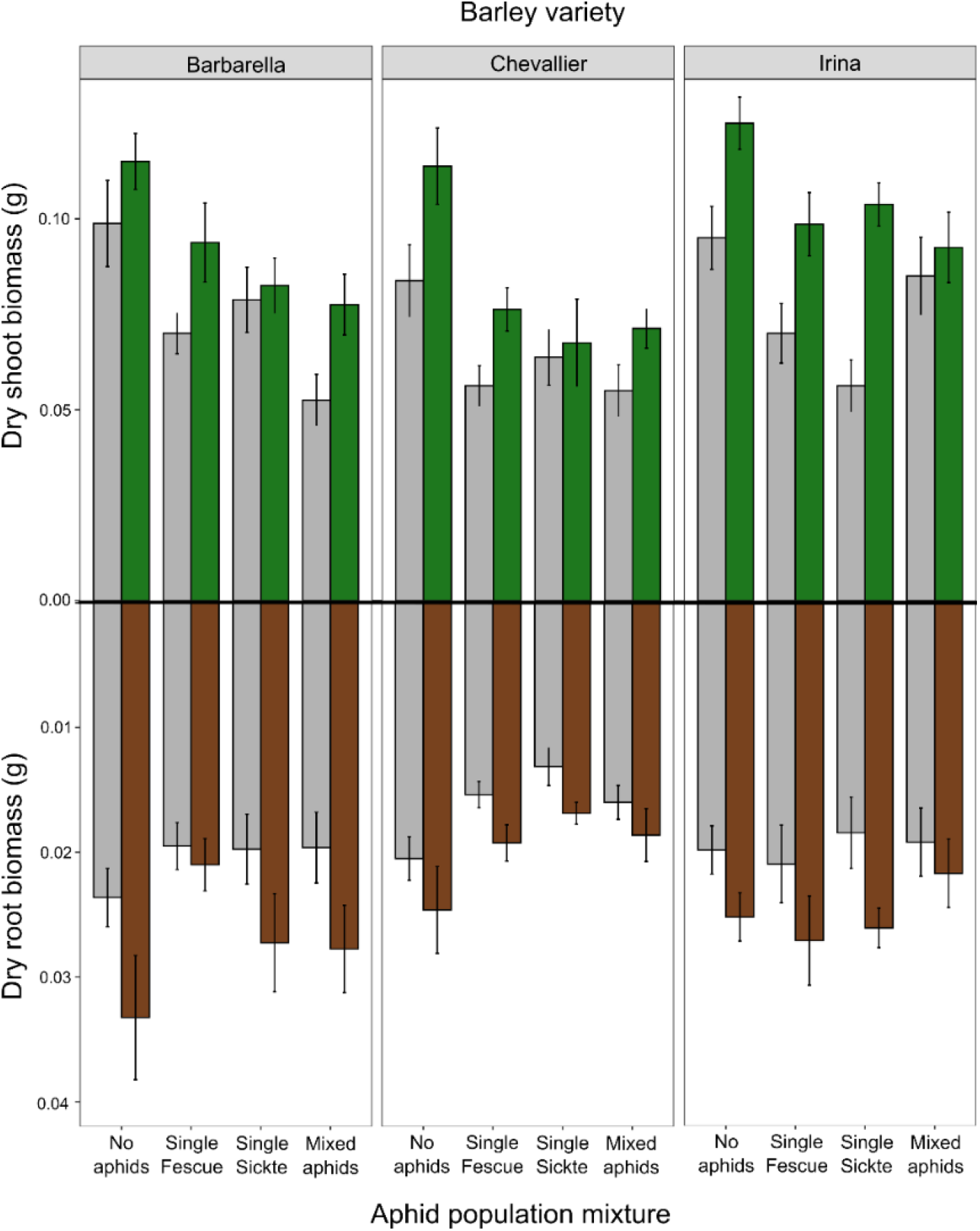
Plant shoot and root biomass responses to bacterial (*Acidovorax radicis*) inoculation across plant variety (Barbarella, Chevallier and Irina) and aphid treatments (control = no aphids; single Fescue, single Sickte = plants infested with one aphid genotype (Fescue or Sickte); and Mixed aphids = plants infested with both aphid genotypes at the same time). Error bars ±1SE, eight replicates per treatment combination.

### Microbe-mediated aphid suppression varies for aphid genotypes leading to rapid evolution in mixed populations

Overall, aphid numbers were reduced on plants inoculated with *A. radicis* at day 14 (F_1,137_=12.46, P<0.001) and day 21 (F_1,135_=13.23, P<0.001), with the *Acidovorax*-suppression effect increasing over time (day x *Acidovorax* F_1,401_=17.14, P<0.001; Fig. 2a). Additionally, at day 21 this effect was dependent on aphid genotype population with a stronger *Acidovorax* suppression effect on aphids in single genotype populations (*Acidovorax* x aphid population F_1,135_=6.01, P=0.015). Both aphid genotypes experienced *Acidovorax*-mediated suppression, with strongest main effects for Sickte aphids (day 21: F_1,89_=38.17, P<0.001; Fig. 2b while for Fescue aphids this was highly dependent on the population mixture where suppression effects were only observed in the single population (*Acidovorax* x aphid population day 21: F_1,87_=11.92, P<0.001; Fig. 2c), with further benefit of feeding in a mixed population on inoculated plants (F_1,87_=5.63, P=0.020; Fig. 2c). In contrast, Sickte aphids experienced higher population growth rates in the single than mixed genotype populations (F_1,89_=5.92, P=0.017; Fig. 2b).

**Figure 2.**
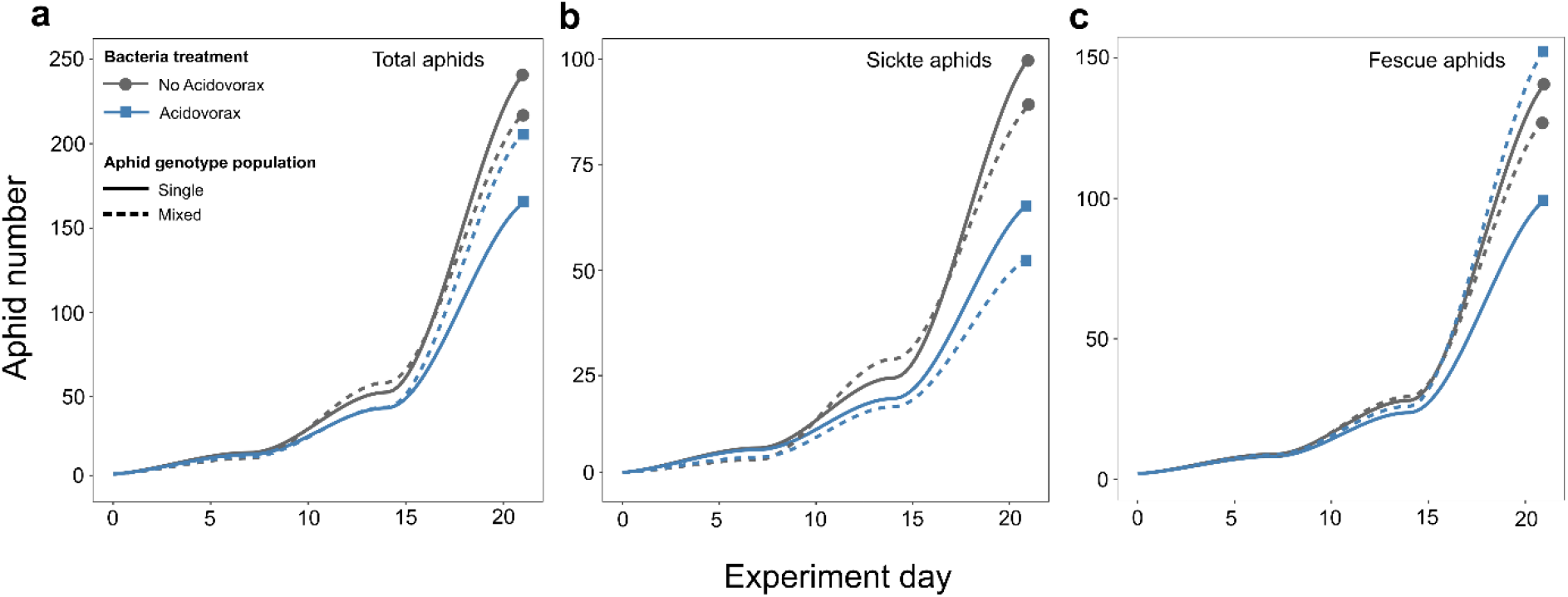
Number of aphids on plants across the experiment (counted on day 7, 14 and 21) separated by *Acidovorax* inoculation treatment (solid line for control plants, dashed lines for inoculated plants) and aphid population mixture (different colours – will change this) for (a) total number of aphids (both genotypes), (b) Sickte aphid genotype and (c) Fescue aphid genotype.

Following, these differential effects of population mixture on the two aphid genotypes led to genotype frequency changes in the mixed populations dependent on *Acidovorax* inoculation (F_1,126_=8.36, P=0.005; Fig. 3a). At day 7, there was no effect of inoculation on aphid genotype frequency, but Fescue aphids had higher growth than Sickte aphids and therefore were more abundant. However, at day 14 there was a more equal number of each aphid genotype on control plants, while on inoculated plants Fescue aphids were more abundant (F_1,38_=4.65, P=0.038). By the end of the experiment at day 21, there was higher abundance of Fescue aphids on all plants compared to Sickte but with a stronger difference on inoculated plants (F_1,38_=12.94, P<0.001) leading to Fescue aphids contributing to almost 75% of the total aphids on inoculated plants (Fig. 3).

**Figure 3.**
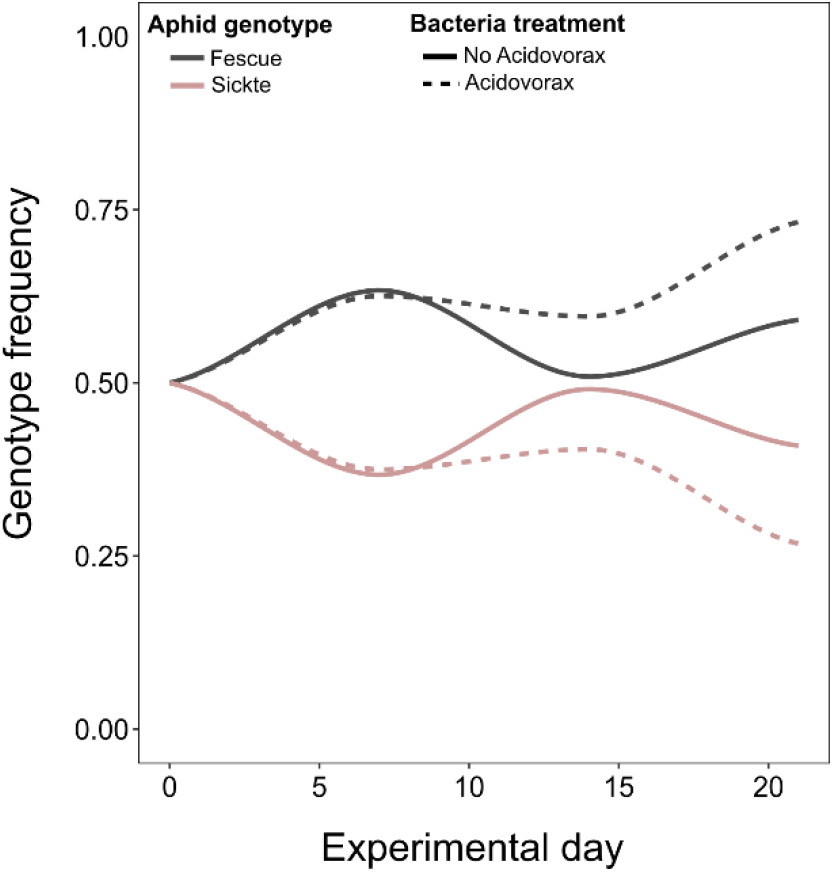
The consequences of differential effects of microbial inoculation and coexistence of aphid genotypes on aphid populations. genotype frequency (from experimental data)

### Aphid population mixture (coexistence) alters microbe-induced plant resistance effects on aphid fecundity and longevity

In the lifetime fecundity and longevity experiment, we found that longevity (days until aphid death, or survival) and aphid lifetime fecundity (total number of offspring produced) were reduced for aphids feeding on inoculated plants (*Acidovorax* main effect: longevity F_1,39_=30.39, P<0.001, Fig. 4a; fecundity F_1,39_=43.54, P<0.001, Fig. 4b). Aphid fecundity (F_1,39_=4.00, P=0.026) and to some extent longevity (F_1,39_=2.81, P=0.071) varied with barley variety, but there were no significant differences among aphid genotypes or population mixtures as main effects (interactions explored in the following).

**Figure 4.**
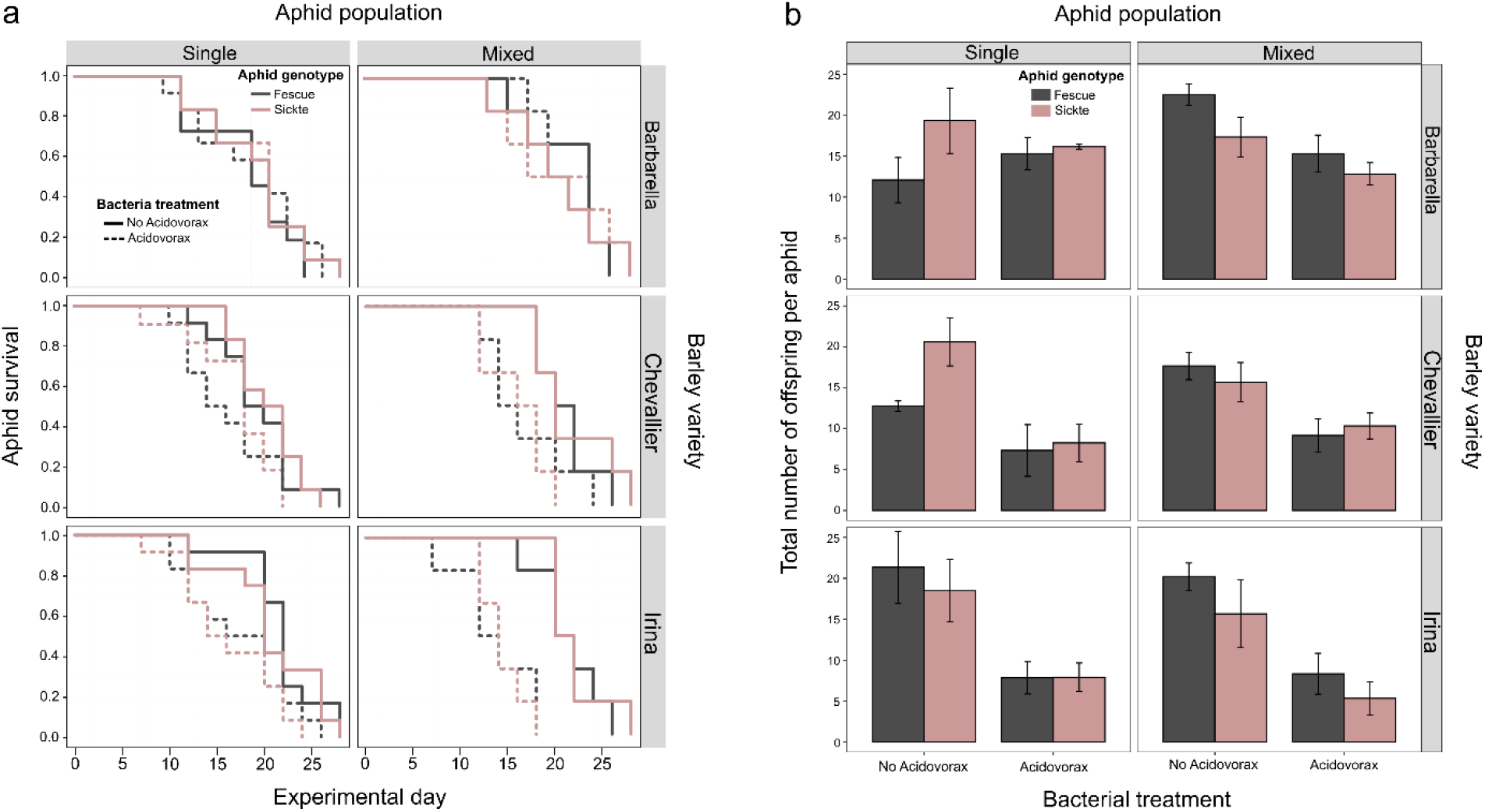
Effect of Acidovorax radicis inoculation of plants on (a) aphid longevity (survival) and (b) aphid lifetime fecundity (total number of offspring per aphid), separated by aphid genotypes within panels, also across plant variety (rows) and aphid population mixture (columns). Error bars ±1SE, six replicates per treatment combination.

The effect of *A. radicis* inoculation on aphid longevity was dependent on the plant variety (*Acidovorax* x barley: F_2,39_=12.03, P<0.001), with reduced survival of aphids on inoculated Chevallier and Irina barley plants and limited effects on Barbarella (Fig 4). No further significant interactions were observed for aphid longevity.

The effect of *A. radicis* inoculation on total aphid fecundity was dependent on the plant variety (*Acidovorax* x barley: F_2,39_=5.80, P=0.006), with a stronger aphid suppression effect on Chevallier and Irina barley plants than Barbarella (Fig 4). Contributing to the reduced total fecundity, aphids feeding on inoculated plants reproduced for fewer days (*Acidovorax* x barley: F_1,39_=12.79, P<0.001), likely in part driven by reduced longevity, but we also showed that surviving aphids were unable to produce as many offspring per day as aphids on control plants (*Acidovorax* main effect: F_1,39_=14.65, P<0.001) (Fig 5). Over the experimental time period, this reduced reproductive output due to *A. radicis* inoculation was also dependent on the aphid population mixture (F_1,39_=8.34, P=0.004), barley variety (F_2,39_=12.74, P=0.002) and the interaction among these three factors (*Acidovorax* x aphid population x barley F_2,39_=6.09, P=0.048) (Fig 5). Further, aphid total fecundity varied as an interaction between aphid genotype and population mixture (Aphid genotype x population: F_1,39_=7.37, P=0.009), where (in agreement with the previous experiment) Fescue aphids experienced higher fecundity in the mixed population while Sickte aphids performed better in the single genotype population. In contrast to the previous experiment, this did not vary across *A. radicis* inoculation treatments.

**Figure 5.**
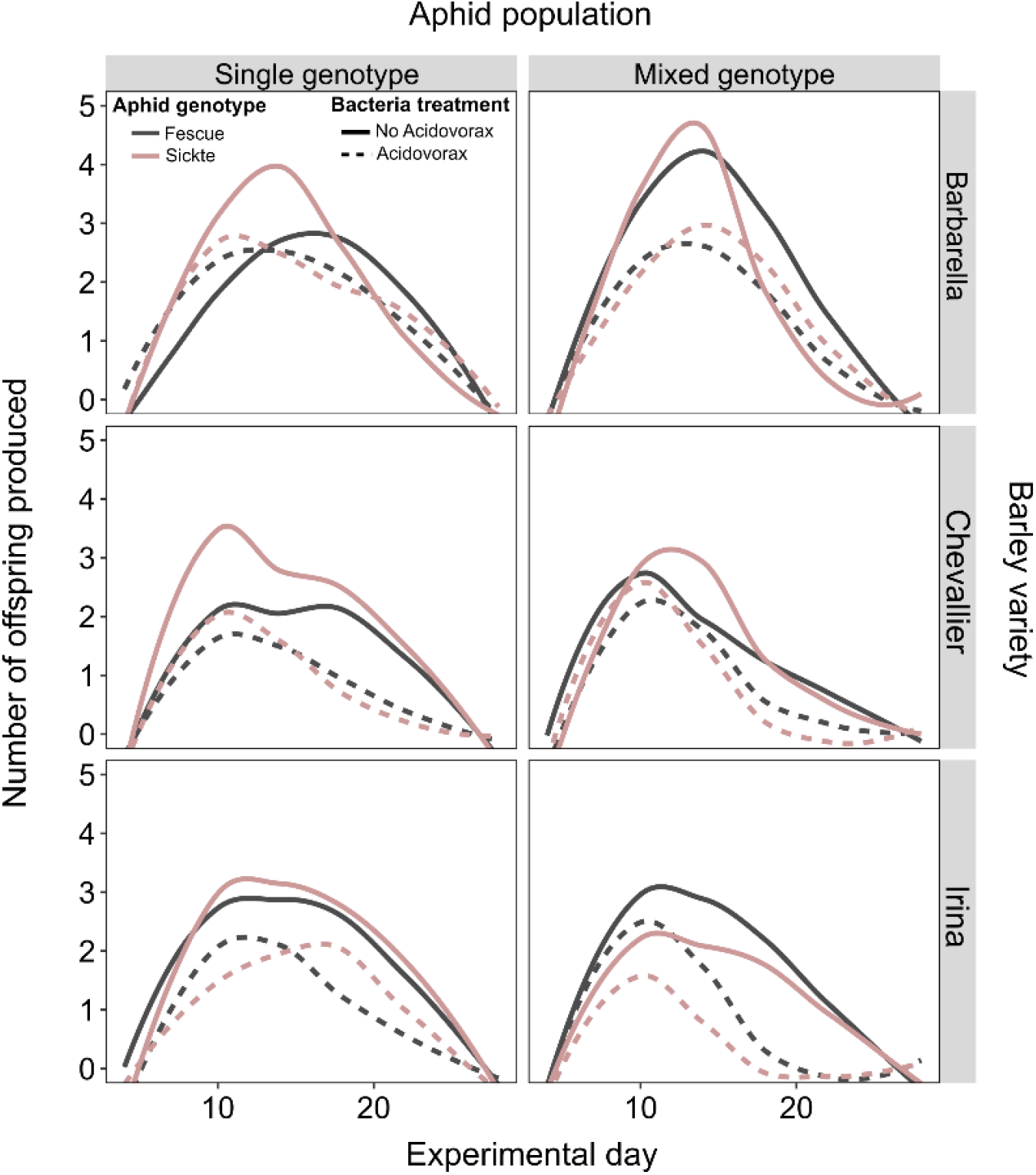
Effect of *Acidovorax radicis* inoculation on offspring production (per surviving aphid) across time (counted every 1-2 days), separated by aphid genotypes within panels, also across plant variety (rows) and aphid population mixture (columns). Six replicates per treatment combination.

## Discussion

Our results show that root-associated plant beneficial microbes have the potential to alter rapid evolution in aphid populations. Overall, plants inoculated with bacteria hosted fewer aphids. This was achieved through reduced survival and daily reproductive output leading to overall lower lifetime fecundity. In mixed populations, the aphids generally performed worse than would be predicted from their performance in single populations. However, one exception was Fescue aphids on inoculated plants where they performed better than would be expected from the single population performance on inoculated plants. In particular, aphid lifetime fecundity, as well as daily reproductive output by surviving aphids, was highly dependent on the population mixture. This means that the aphids reared as a single genotype population performed differently to those reared in mixed genotype populations leading to changes in genotype frequency (rapid evolution; Turcotte, Reznick and Hare (2013)). We also showed that aphid suppression and plant-growth promotion effects varied across barley genotypes dependent on the aphid genotype and populations highlighting that plant genetic variation, as well as aphid genetic variation, underlies many of these responses to the interactions among the community members.

Susceptibility to aphid herbivory varied among the three plant varieties as did their response to *A. radici*s inoculation. Aphids decreased aboveground plant biomass of all varieties while *A. radici*s increased it, and the outcome of this interaction varied among plant varieties; suggesting that interactions between aphids and soil bacteria could potentially drive plant evolution (Kempel *et al*. 2013). We found that these interactions affecting the plant’s response to bacterial inoculation were primarily driven by aphid presence/absence or the single versus mixed populations, rather than aphid genotypes. Thus, we suggest that this is mediated by bacteria-induced plant physiological changes that alter the aphid’s performance, e.g. plant defensive phenotypes (Sugio *et al*. 2015; Schädler & Ballhorn 2016), which can also feedback to alter aphid communities in a density-dependent manner. Previously, we have found larger plants to host more aphids in this system (Zytynska *et al*. 2020), but here for inoculated plants we found this relationship to differ across plant cultivars (positive for Barbarella but negative for Irina plants, where larger plants had fewer aphids). Thus, effects of bacterial inoculation not only affected the plant growth or aphid numbers but altered interactions among these. This could be explained by the plant trade-off between allocating energy to growth or anti-herbivory defence (Züst & Agrawal 2017). Interactions with the inoculated bacteria that simultaneously increases plant growth and plant defence against aphids could eliminate this trade-off as demonstrated in a recent paper with *Epichloë* fungal endophytes (Bastías, Gianoli & Gundel 2021). In our experiment, inoculated Irina plants experienced both high growth promotion and aphid suppression, while inoculated Barbarella plants experienced average growth promotion and lower aphid suppression. It may be concluded that plant genetic variation in the response to bacterial inoculation mediates the effect on this growth-defence trade-off, dependent on aphid genotype diversity (Stam *et al*. 2014).

The effects of plant bacterial inoculation on aphid populations shifted the aphid genotype frequency and led to rapid evolution of the mixed populations. This was driven by reproductive output (fecundity) rather than differential survival of the aphid genotypes and was dependent on the genotype population mixture. Since the temperature in the experiment was above the threshold temperature that male aphids are produced, the changes in the relative genotype frequency in aphid population are due to the differentially growth rate of the two clonally reproducing genotypes. Different aphid genotypes induce variable plant defences when feeding, with fitness consequences for other aphid genotypes (Zytynska *et al*. 2016), and these will also vary among plant varieties (Delp *et al*. 2009). Aphids also release salivary proteins into the plant to facilitate feeding and suppress plant defences (Yates & Michel 2018) and it is likely that genetic variation in both the aphid and plant influence the outcome of these interactions. We found that Fescue aphids benefited from feeding in the mixed populations, particularly in the first experiment where aphids remained on the plant, while Sickte aphids benefited more in the single populations. One hypothesis to be tested further is that Sickte aphids release effectors that also facilitate Fescue aphid feeding, while the Fescue aphids do not reciprocate. We have also shown that soil bacteria alter the outcome of these plant-aphid interactions, and it is possible that this occurs through altering the ability of the aphids to alter local plant physiology to their benefit; however, this also remains to be tested.

Intra-species variation has been demonstrated to have great impact on species coexistence (Hart, Turcotte & Levine 2019), consumer-resource dynamics (Raffard *et al*. 2021) and receives increasing attention in biodiversity-ecosystem functioning studies (Violle *et al*. 2012; Whitham *et al*. 2020). Our experiment indicated that mixed aphid genotype populations resulted in variable shoot damage across plant varieties compared to the single genotype treatment dependent on soil bacteria inoculation, and interactions among the aphid-suppressing bacteria treatment and between aphid genotypes altered aphid growth rates with consequences for aphid rapid evolution in the mixed populations. Several studies have demonstrated that agricultural practices, like crops breeding, could trigger pest evolution (Turcotte, Reznick & Daniel Hare 2013; Turcotte *et al*. 2015; Züst & Agrawal 2016; Simon & Peccoud 2018). Our studies demonstrated that such effects could be altered by interactions with soil bacteria and suggests that soil microbes may shift intra-species diversity-ecosystem functioning relationships.

## Acknowledgements

This work was supported by National Science Foundation of China (31870417) grant to XX and a BBSRC (UKRI) David Phillips Fellowship BB/S010556/1 to SEZ.

